# X chromosome dosage drives statin-induced dysglycemia and mitochondrial dysfunction

**DOI:** 10.1101/2022.08.29.505759

**Authors:** Peixiang Zhang, Joseph J. Munier, Laurent Vergnes, Carrie B. Wiese, Jenny C. Link, Fahim Abbasi, Emilio Ronquillo, Antonio Muñoz, Yu-Lin Kuang, Meng Liu, Gabriela Sanchez, Akinyemi Oni-Orisan, Carlos Iribarren, Michael J. McPhaul, Daniel K. Nomura, Joshua W. Knowles, Ronald M. Krauss, Marisa W. Medina, Karen Reue

**Affiliations:** Human Genetics, David Geffen School of Medicine, University of California, Los Angeles, CA 90095; Molecular, Cellular & Integrative Physiology, University of California, Los Angeles, CA; Molecular Biology Institute, University of California, Los Angeles, CA; Division of Cardiovascular Medicine and Cardiovascular Institute, Diabetes Research Center, Stanford University School of Medicine, Stanford, CA; School of Medicine, University of California, San Francisco, Oakland, CA; Division of Research, Kaiser Permanente, Oakland, CA; Institute for Human Genetics, University of California, San Francisco, CA; Quest Diagnostics Nichols Institute, San Juan Capistrano, CA 92675; Nutritional Sciences and Toxicology, and Novartis-Berkeley Center of Proteomics and Chemistry Technologies, University of California, Berkeley, Berkeley, CA

## Abstract

Statin drugs lower blood cholesterol levels for cardiovascular disease prevention. Women are more likely than men to experience adverse statin effects, particularly new-onset diabetes (NOD) and muscle weakness. We determined that female mice are more susceptible than males to glucose intolerance, fasting hyperglycemia, and muscle weakness after short-term statin treatment. Lipidomic, transcriptomic, and biochemical analyses identified reduced docosahexaenoic acid (DHA) levels, and impaired redox tone and mitochondrial respiration specifically in statin-treated female mice. Statin adverse effects could be prevented in females by complementation with a source of DHA. Statin adverse effects segregated with XX chromosome complement, and specifically dosage of the *Kdm5c* gene, which regulates fatty acid gene expression and has differential expression levels in females and males. In humans, we found that women experience more severe reductions than men in DHA levels after short-term statin administration, and that DHA reduction was correlated with increases in fasting glucose levels. Furthermore, induced pluripotent stem cells derived from women, but not men, who developed NOD exhibited impaired mitochondrial function when treated with statin. Overall, our studies identify biochemical mechanisms, biomarkers, and a genetic risk factor for susceptibility to statin adverse effects, and point to DHA supplementation as a preventive co-therapy.

## Introduction

Statins, or HMG-CoA reductase enzyme inhibitors, are the most widely prescribed drug class for reducing blood cholesterol levels and risk of cardiovascular disease (Mach et al., 2018). However, statins may have adverse effects, the most common of which are myopathy (Thompson et al., 2016) and new-onset diabetes (Carter et al., 2013; Danaei et al., 2013; Navarese et al., 2013; Preiss and Sattar, 2011; Sattar et al., 2010), and there is evidence that both are significantly more common in women than men. Myopathy symptoms range from minor pain and weakness (in 5–10% of users) to life-threatening rhabdomyolysis (in <0.1% of users) (Thompson et al., 2016). A meta-analysis indicated that female sex increases the odds of stain-associated myopathy by nearly 2-fold, although differences exist across studies (Nguyen et al., 2018). Analyses of incident diabetes in statin clinical trials with stratification by sex have shown greater diabetes risk in women compared to men (Goodarzi et al., 2013; Mora et al., 2010).

We exploited the sex differences in susceptibility to statin-associated myopathy and new-onset diabetes (NOD) to identify mechanisms that contribute to these adverse effects. We used mouse models that allow comparison of males and females on the same genetic background, as well as analysis of the contribution of gonadal sex differences (ovaries vs. testes) and chromosomal sex differences (XX vs. XY), to identify molecular and genetic determinants of statin-related impairment of glucose homeostasis and muscle strength.

We determined that statin drug interacts with female sex, particularly XX sex chromosome complement, to alter the lipidome and transcriptome in ways that impact mitochondrial function and cellular redox tone. We further identified nutritional and genetic manipulations that prevent statin adverse effects in female mice. Ultimately, we verified the findings from mice in human cohorts, showing differential alterations in lipid metabolism and mitochondrial function in women compared to men treated with statin. Our findings suggest a potential nutritional co-therapy to statins that may prevent adverse effects in vulnerable individuals.

## Results

### Female sex promotes statin-induced glucose intolerance and reduced muscle strength in the mouse

We treated cohorts of male and female C57BL/6J mice (aged 10 weeks) with a chow diet or the same chow diet containing simvastatin (dose equivalent to human dose of 80 mg/day). Independent mouse cohorts were assessed for fasting glucose and glucose tolerance at 2, 4, 8, and 16 weeks of statin treatment. Compared to their control counterparts, female mice treated with statin for as little as 2 weeks exhibited impaired glucose tolerance (as assessed by area-under-the-curve for glucose tolerance tests) (**Fig. 1A**). Glucose intolerance persisted in female cohorts treated with statin for 4, 8, and 16 weeks (**Fig. 1A**). Statin-treated male mice also experienced impaired glucose tolerance, but this was delayed relative to female mice, occurring only after 8 or 16 weeks of statin treatment (**Fig. 1B**). Furthermore, female mice developed fasting hyperglycemia, which was evident at 4, 8, and 16 weeks after statin treatment (**Fig. 1C)** whereas male mice did not alter fasting glucose levels in response to statin **(Fig. 1D)**. Statin treatment did not alter insulin levels in male or female mice, but male mice had higher absolute insulin levels compared to females with either chow alone or chow containing statin (**Suppl. Fig. 1**).

**Figure 1.**
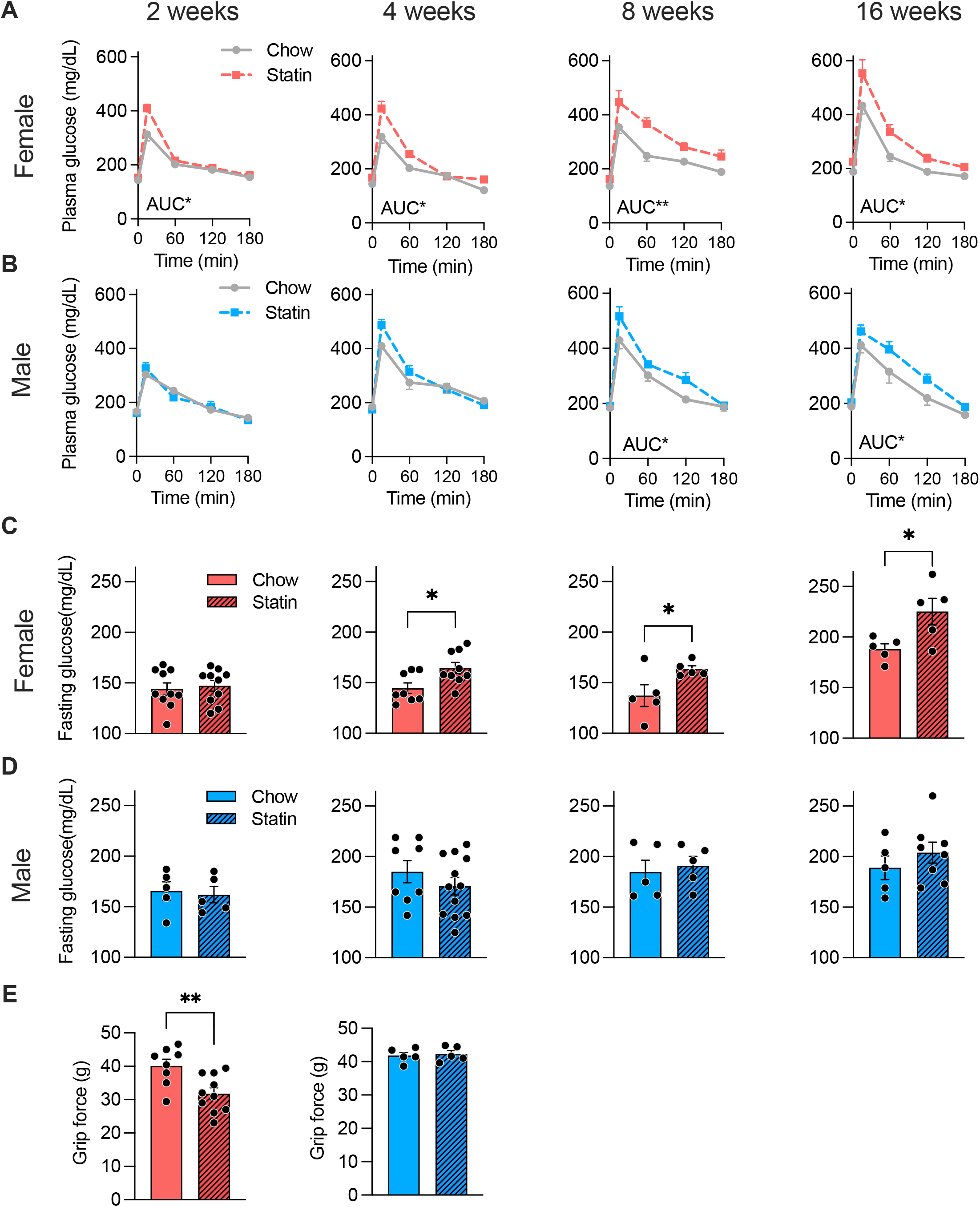
Female sex exacerbates statin-induced changes in glucose homeostasis and grip strength in mouse. Independent cohorts of C57BL/6J mice (n=5−12 per sex and treatment) were fed chow or chow containing simvastatin (0.1 g/kg) for 2, 4, 8, or 16 weeks. **(A–B)** Glucose levels during a glucose tolerance test after indicated time on statin treatment in female and male mice. AUC, area under the curve. **(C–D)** Fasting glucose levels after indicated times on statin treatment in female (red) and male (blue) mice. **(E)** Forelimb grip strength after 8 weeks statin treatment in female and male mice. Grip force is measured in grams (g). Values represent mean ± SEM. *, *p* < 0.05; **, *p* < 0.01, statin vs. chow, via unpaired *t*-test. See also Figure S1.

To test for statin effects on muscle strength, we measured grip strength in the forelimbs with a grip bar strength dynamometer. Male mice had similar grip strength force on chow and statin, but female mice experienced a 20% reduction in grip strength after 8 weeks of statin treatment (**Fig. 1E**).

### Statin adverse effects in female mice are associated with reduced ω-3 fatty acid levels

Most statins, including simvastatin, are metabolized in the liver. Statins reduce hepatic cholesterol synthesis by inhibiting the rate-limiting enzyme in the mevalonate pathway. This has the potential to alter the levels of lipid intermediates downstream of mevalonate, including cholesterol precursors, isoprenoids, and Coenzyme Q10 (CoQ10). We profiled 130 metabolite species in liver from male and female mice on control and statin diets for 4 weeks, a point at which only female mice developed impaired glucose tolerance (see Fig. 1). We assessed metabolite levels after statin treatment compared to levels on the control diet for each sex.

Females treated with statin experienced more widespread changes in hepatic lipidome and transcriptome compared to males (**Fig. 2A**,**B**). In particular, statin-treated females showed reduction in levels of several lipid species (**Fig. 2A**; **Table S1**). With regard to lipid species within the mevalonate pathway, statin treatment reduced cholesterol in both sexes (females showed a significant reduction in free cholesterol, and males in cholesterol esters) (Suppl. **Fig. 2A**). Males trended to reduced CoQ10 levels after statin treatment. Statin treatment in females reduced geranylgeranyl pyrophosphate levels by ∼30%, whereas males showed a trend to reduced levels of farnesyl pyrophosphate. Neither skeletal muscle nor plasma from either sex showed alterations in cholesterol biosynthetic pathway components in response to statins (Table S1).

**Figure 2.**
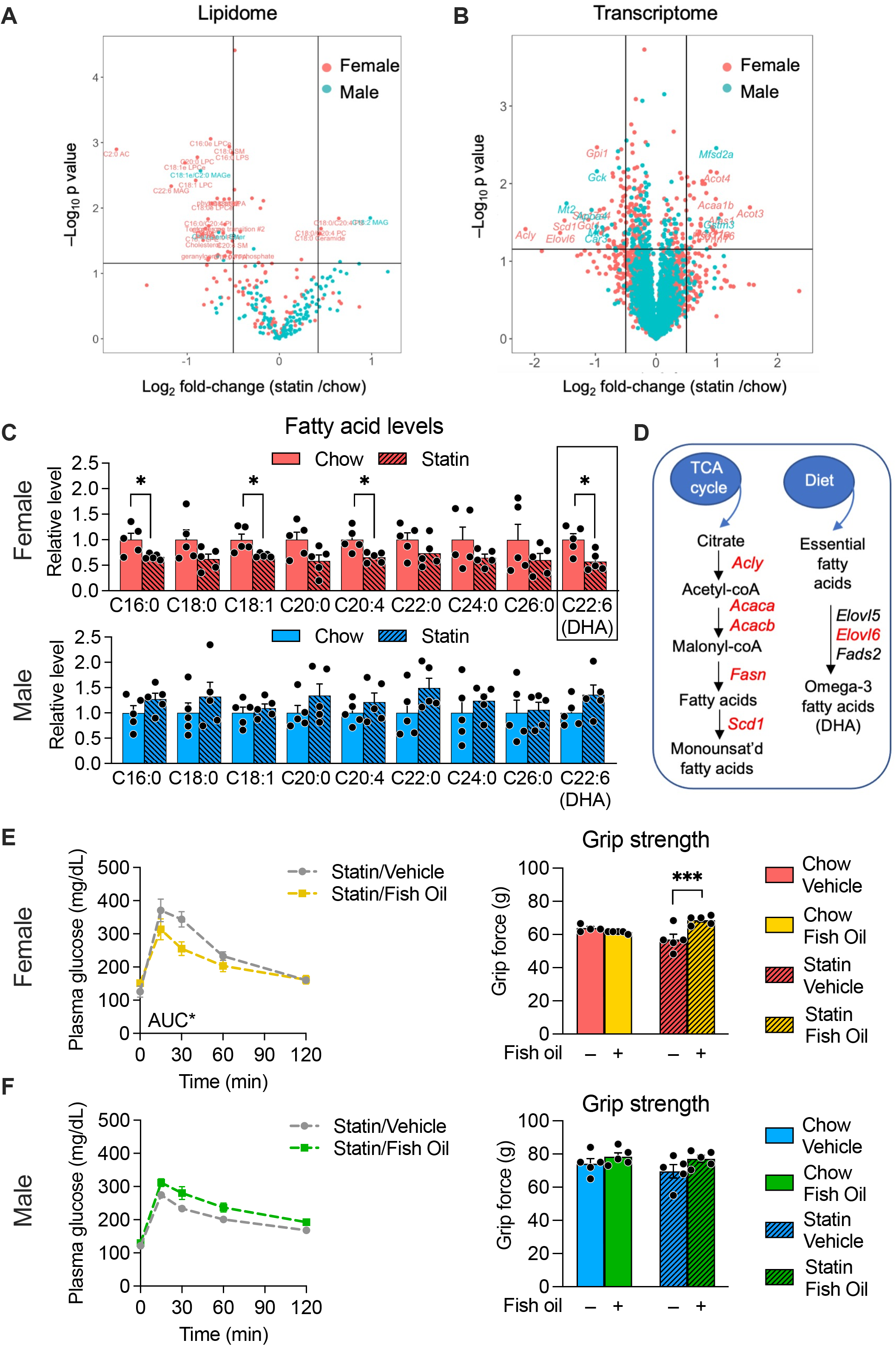
Female-specific statin-induced reductions in fatty acids, and prevention of adverse effects with fatty acid complementation in mouse. **(A)** Mass spectrometric profiling of liver lipidome in mice treated with statin for 4 weeks (n=5 per sex and treatment). Lipid species with significant differences between statin and control diets within each sex (p < 0.05) are shown above the horizontal line. The vertical lines delineate a 50% decrease (to the left) or increase (to the right) in levels in response to statin. **(B)** Volcano plot of RNA-seq gene expression data from liver of mice in (A). Genes with significant differences between statin and control diets within each sex (p < 0.05 with correction for multiple testing) are shown above the horizontal line. The vertical lines delineate 0.5-fold decrease (to the left) or increase (to the right) in mRNA levels in response to statin. **(C)** Relative levels of fatty acid species in females and males in mice fed chow without or with statin. Fatty acids are represented as number of carbons followed by number of double bonds. DHA, docosahexaenoic acid. **(D)** Pathways for synthesis of monounsaturated (*left*) and polyunsaturated fatty acids (*right*). Gene names in red exhibit reduced expression in female mice treated with statin. **(E**,**F)** Mice were fed chow or statin-containing diet with daily administration of either fish oil (as a source of ω-3 fatty acids) or vehicle. After 5 weeks, glucose tolerance and grip strength were measured in (E) female or (F) male mice. N = 5/sex on each treatment. AUC, area under the curve. Error bars represent SEM. Data for each gene, metabolite, or mitochondrial complex were analyzed by 2-way ANOVA, and if significant at p < 0.05, post-hoc analysis was by pairwise t-tests for comparisons indicated by brackets (*, *p* < 0.05; ***, *p* < 0.001 for pair-wise tests).

Beyond mevalonate pathway lipid species, statin treatment reduced hepatic levels of several fatty acid and phospholipid species exclusively in females. These included a 30% reduction medium- and long-chain fatty acid species (C16:0, C18:1, and C20:4) and a 40% reduction in the essential ω-3 polyunsaturated fatty acid, docosahexaenoic acid (DHA, C22:6) (**Fig. 2C** and Table S1); the ω-3 fatty acid eicosapentaenoic acid (EPA, C20:5) was not detectable in our assay. We also detected female-specific reductions in levels of sphingomyelins (C18:0, C18:1, C20:4) and increased levels of ceramides (C18:0) (Table S1), which are derived from sphingomyelin hydrolysis.

We detected sex differences in the liver transcriptome that correlate with the lipidome. Consistent with the reduced hepatic fatty acid levels in females, statin treatment led to reduced expression of genes required for fatty acid synthesis (*Acly, Acaca, Acacb, Fasn*), for desaturation of medium-chain fatty acids (*Scd1*), and for synthesis of ω-3 fatty acids (*Elovl6*) compared to their expression levels on control diet (**Fig. 2D, Suppl. Fig. 2B**).

Reduced ω-3 fatty acid levels have previously been associated with glucose intolerance and insulin resistance (Abbott et al., 2016; Eguchi et al., 2011; Flachs et al., 2014; Lamping et al., 2013; Lepretti et al., 2018; O’Mahoney et al., 2018; Ramel et al., 2008). We therefore hypothesized that overcoming the deficit in DHA levels might prevent the adverse physiological effects that we observed in statin-treated female mice. To test this hypothesis, we fed male and female mice the control or statin-containing diets and provided either fish oil (a source of ω-3 fatty acids, DHA and EPA) in coconut oil vehicle, or coconut oil vehicle alone (which contains primarily saturated fatty acids). Fish/coconut oil dosing was performed by oral gavage 5 times/week. After 5 weeks, glucose tolerance and grip strength were determined. Fish oil co-therapy with statin prevented glucose intolerance and reduction in grip strength in females, with no significant effects on these traits in males (**Fig. 2E**,**F**).

### Statin treatment impairs the DHA–Nrf2–glutathione axis and mitochondrial activity in female mice

DHA protects against oxidative stress through induction of the glutathione anti-oxidant system (Arab et al., 2006; Drolet et al., 2021; Patten et al., 2013; Zgórzyńska et al., 2017). The female-specific reduction in DHA levels led us to investigate whether the glutathione axis is compromised in statin-treated females. Glutathione (GSH) is a tripeptide composed of glutamate, cysteine, and glycine that has a critical role in protecting against environmental toxins, including drugs (Chen et al., 2013). DHA and other polyunsaturated fatty acids activate the redox-sensitive transcription factor, Nrf2, which promotes GSH synthesis through expression of *Gclc* (encoding the rate-limiting enzyme in GSH synthesis), as well as other genes involved in maintenance of cellular redox state (**Fig. 3A**) (Abrescia et al., 2020; Mani et al., 2013; Zgórzyńska et al., 2017).

**Figure 3.**
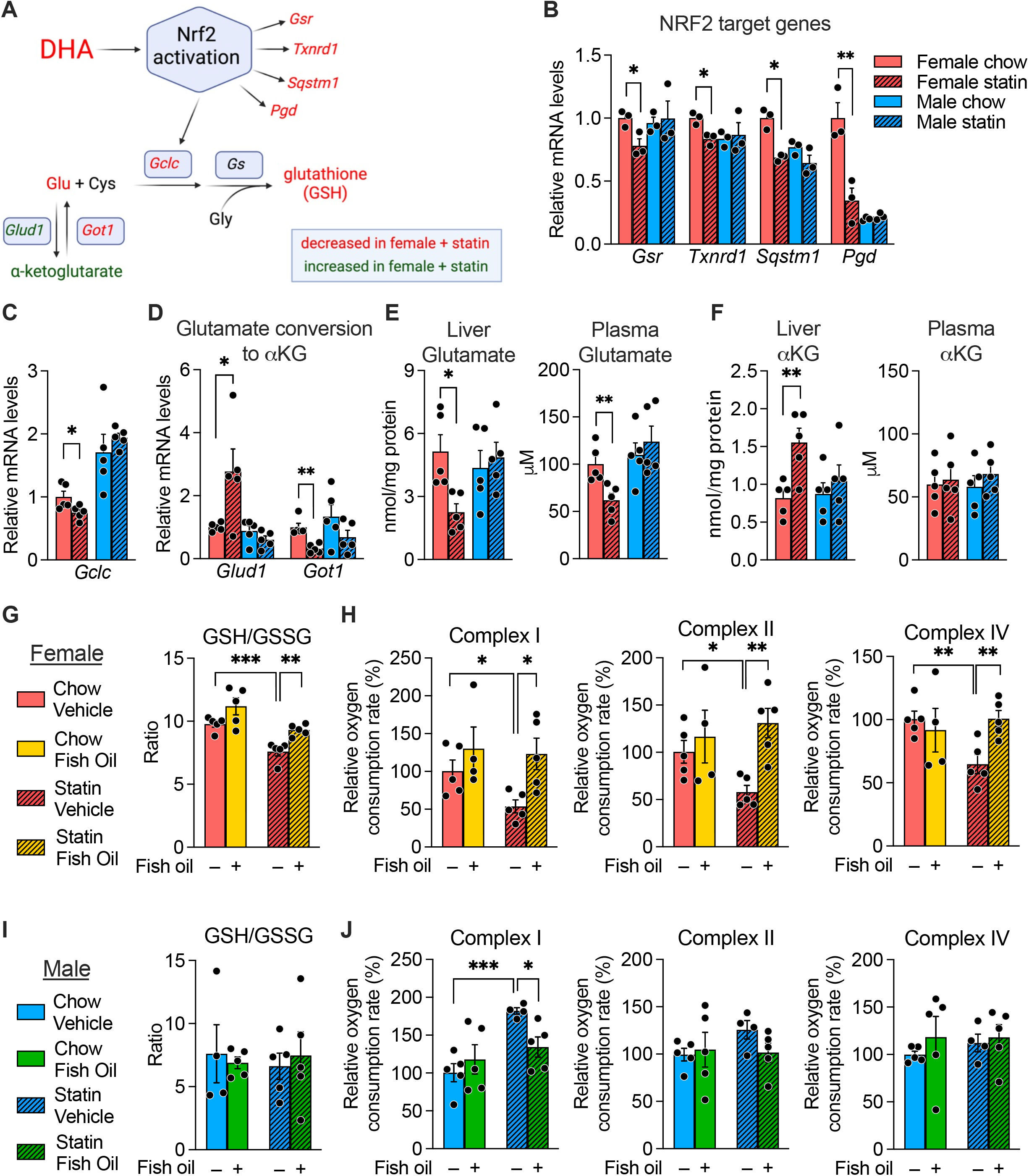
Statin impairs, and fish oil normalizes, glutathione levels and mitochondrial respiration in female mice. **(A)** DHA promotes production of the anti-oxidant, glutathione, through activation of Nrf2 and regulation of its transcriptional targets, including the gene encoding the rate-limiting enzyme in glutathione biosynthesis, *Gclc*. Red text, mRNA and metabolite levels that are decreased in females in response to statin; green text, levels that are increased in females in response to statin; black text, not measured. Data for components shown in red and green are provided in subsequent panels of the figure; DHA levels are shown in Fig. 2C. **(B**,**C)** Hepatic mRNA expression levels for glutathione synthetic gene, *Gclc*, and additional Nrf2 target genes (*Gsr, Txnrd1, Pgd, Sqstm1*). **(D)** Expression levels for *Glud1* and *Got1*, which regulate glutamate (Glu) and α-ketoglutarate interconversion. (**E)** Glutamate and **(F)** α-ketoglutarate levels in liver and plasma. **(G–J)** Effect of statin treatment in combination with fish oil or vehicle on glutathione levels (ratio of reduced to oxidized forms) in female (G) and male mice (I), and on mitochondrial respiration of complexes I, II and IV in female (H) and male mice (J).

Consistent with reduced Nrf2 activation, statin-treated females had reduced expression levels of Nrf2 target genes including *Gsr* (glutathione reductase), *Txnrd1* (thioredoxin reductase) *Pgd* (phosphogluconate dehydrogenase), *Sqstm1* (sequestosome 1), and importantly, the rate-limiting enzyme in glutathione synthesis, *Gclc* (**Fig. 3B**,**C**). Furthermore, statin-treated females experienced a 50% reduction in the levels of the glutathione precursor, glutamate. Gene expression alterations in *Glud1* and *Got1* are consistent with a shift in equilibrium to reduce glutamate levels at the expense of increased α-ketoglutarate (**Fig. 3A**,**D**). Indeed, liver and plasma levels of glutamate were reduced in statin-treated females (**Fig. 3E**), whereas liver levels of α-ketoglutarate were increased (**Fig. 3F**). Collectively, these findings indicate that statin reduces activation of the DHA–Nrf2–glutathione axis in females.

The reduced form of glutathione (GSH) is a critical antioxidant for maintenance of the mitochondrial redox environment (Marí et al., 2009). We therefore tested whether mitochondrial activity was also influenced by statin treatment. Statin treatment of female mice led to a decrease in hepatic GSH/GSSG ratio (a measure of reduced to oxidized forms of glutathione), and reductions in mitochondrial complex I, II and IV activity (**Fig. 3G**,**H, red bars**). When females received fish oil in combination with statin treatment, the decrements in GSH levels and mitochondrial respiration were largely prevented (**Fig. 3G**,**H, yellow bars**). Male mice did not experience reductions in GSH/GSSG levels nor mitochondrial activity with statin (**Fig. 3I**,**J**), but rather displayed an increase in complex I activity on statin, which was normalized by fish oil co-therapy.

### XX chromosomal sex and *Kdm5c* gene dosage increase risk for statin adverse effects

We next investigated what determinants of biological sex contribute to enhanced susceptibility to adverse statin effects in females. Sex differences between males and females can be parsed into genetic sex (XX vs. XY chromosome complement), which is a source of gene dosage differences between males and females, and gonadal sex (ovaries vs. testes), which confers differential gonadal hormone levels in females and males. We used the Four Core Genotypes mouse model to interrogate genetic and gonadal sex contributions. This model consists of four sex genotypes—mice with XX chromosomes and either female or male gonads, and mice with XY chromosomes and either female or male gonads (Mauvais-Jarvis et al., 2017) (**Fig. 4A**).

**Figure 4.**
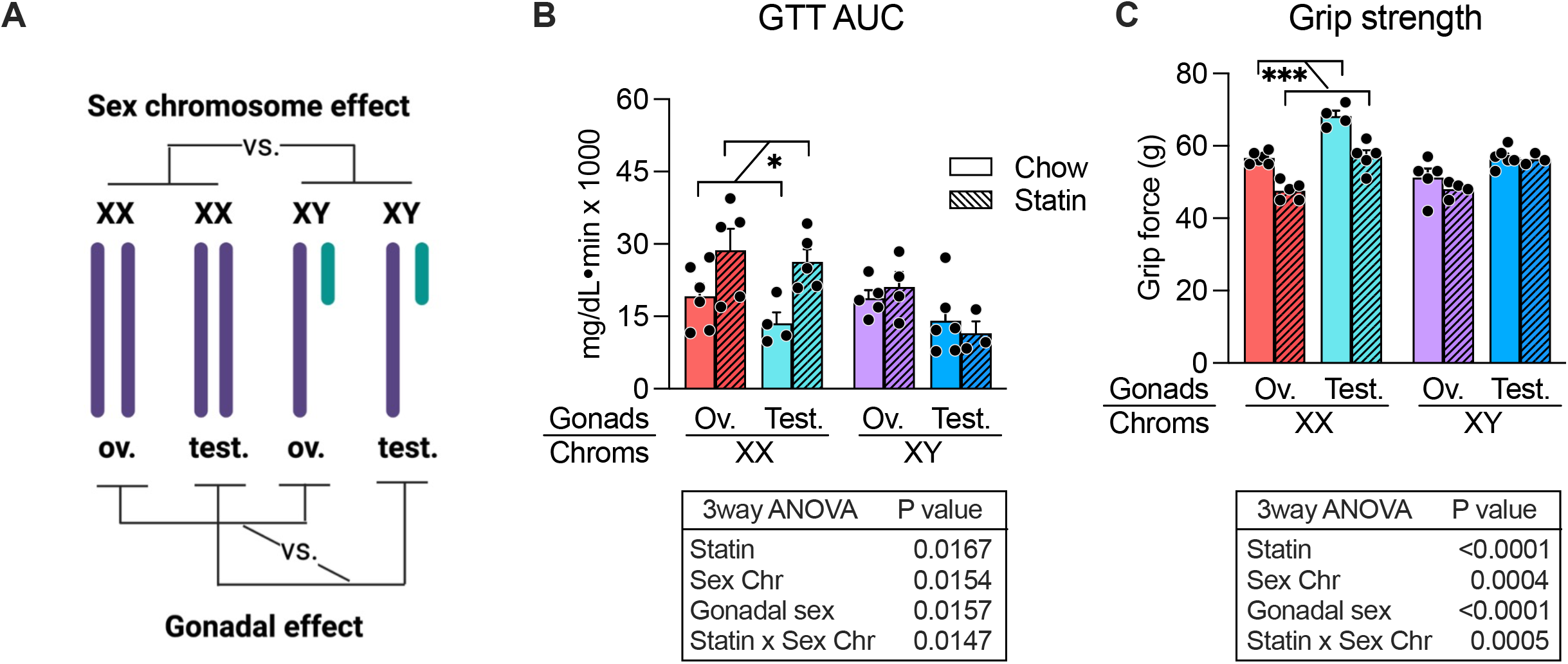
XX chromosome complement segregates with statin adverse effects in mouse. **(A)** The Four Core Genotypes mouse model was used to assess the relative roles of sex chromosomes and gonadal sex in statin adverse effects. The four genotypes of this model are shown with the comparisons that allow assessment of chromosome vs. gonadal effects. Ov., ovaries; Test., testes. **(B)** Glucose tolerance test AUC (area under the curve) after 6 weeks on statin. **(C)** Grip strength after 7 weeks on statin. Values represent mean ± SEM. Results were analyzed by 3-way ANOVA using Bonferroni correction (α = 0.0125); if significant at p<0.05, post-hoc analysis was by pairwise t-tests for comparisons indicated by brackets (*, *p* < 0.05; **, p < 0.01; ***, *p* < 0.001 for pair-wise tests).

We evaluated statin adverse effects in C57BL/6 Four Core Genotypes mice fed control or statin diet for 8 weeks. We analyzed these data by 3-way ANOVA using gonadal sex, chromosomal sex, and statin treatment as variables. We detected significant effects of sex chromosome complement on statin-induced impairment in glucose tolerance and grip strength. After statin treatment, mice with XX chromosomes having either testes or ovaries showed impaired glucose tolerance (increased glucose tolerance test area under the curve (**Fig. 4B, red and teal bars**); by contrast, XY mice were protected from statin-induced changes in glucose tolerance (**Fig. 4B, purple and blue bars**). XX mice also experienced reduced grip strength with statin treatment, whereas XY mice did not alter grip strength in response to statin (**Fig. 4C**). Thus, XX chromosome complement is a risk factor for statin adverse effects.

XX chromosome complement may influence metabolism through genes that escape X-chromosome inactivation and are therefore expressed at higher levels in XX compared to XY cells (Davis et al., 2020; Itoh et al., 2019; Link et al., 2020). We previously screened known X-escape genes for their sex-biased expression in mouse liver and identified the four genes that showed greatest differences in hepatic expression levels between XX and XY mice (Chen et al., 2012; Link et al., 2020). We noted that one of these genes with higher expression levels in female mice, *Kdm5c*, encodes a histone demethylase that represses fatty acid biosynthetic gene expression, including fatty acid synthase (*Fasn*) (Zhang et al., 2020), which are reduced by statin exclusively in female mice (see Fig. D). Furthermore, KDM5C histone demethylase activity requires α-ketoglutarate as a co-factor (Iwase et al., 2007), and α-ketoglutarate levels are increased in statin-treated females (see Fig. 3F).

We hypothesized that the higher *Kdm5c* expression levels in XX compared to XY individuals (Link et al., 2020) may contribute to sex differences in fatty acid biosynthetic gene expression, redox tone, and statin adverse effects. To test this, we generated female mice with either the standard *Kdm5c* gene dosage (two active *Kdm5c* alleles, *Kdm5c*^+/+^) or the *Kdm5c* gene dosage present in male mice (one active *Kdm5c* allele, *Kdm5c*^+/–^) (**Fig. 5A**); female *Kdm5c*^+/–^ mice have ∼50% of the KDM5C protein levels present in female *Kdm5c*^+/+^ mice (Link et al., 2020). When treated with statin, *Kdm5c*^+/+^ mice were susceptible to statin-induced glucose intolerance as we have seen throughout our studies with female mice, but *Kdm5c*^+/–^ mice were protected (**Fig. 5B**). Grip strength, however, was not significantly impacted by statin treatment in mice with either *Kdm5c* genotype. *Kdm5c*^+/–^ mice also were protected from statin-induced reduction in GSH/GSSG ratio (**Fig. 5C**) and mitochondrial respiration (**Fig. 5D**), and resisted statin-induced reduction of fatty acid gene expression levels that occur in *Kdm5c*^+/+^ mice (**Fig. 5E**). Thus, altering female *Kdm5c* gene dosage to that normally present in males largely prevents statin-induced impairment of glucose tolerance, redox tone, mitochondrial activity, and fatty acid gene expression.

**Figure 5.**
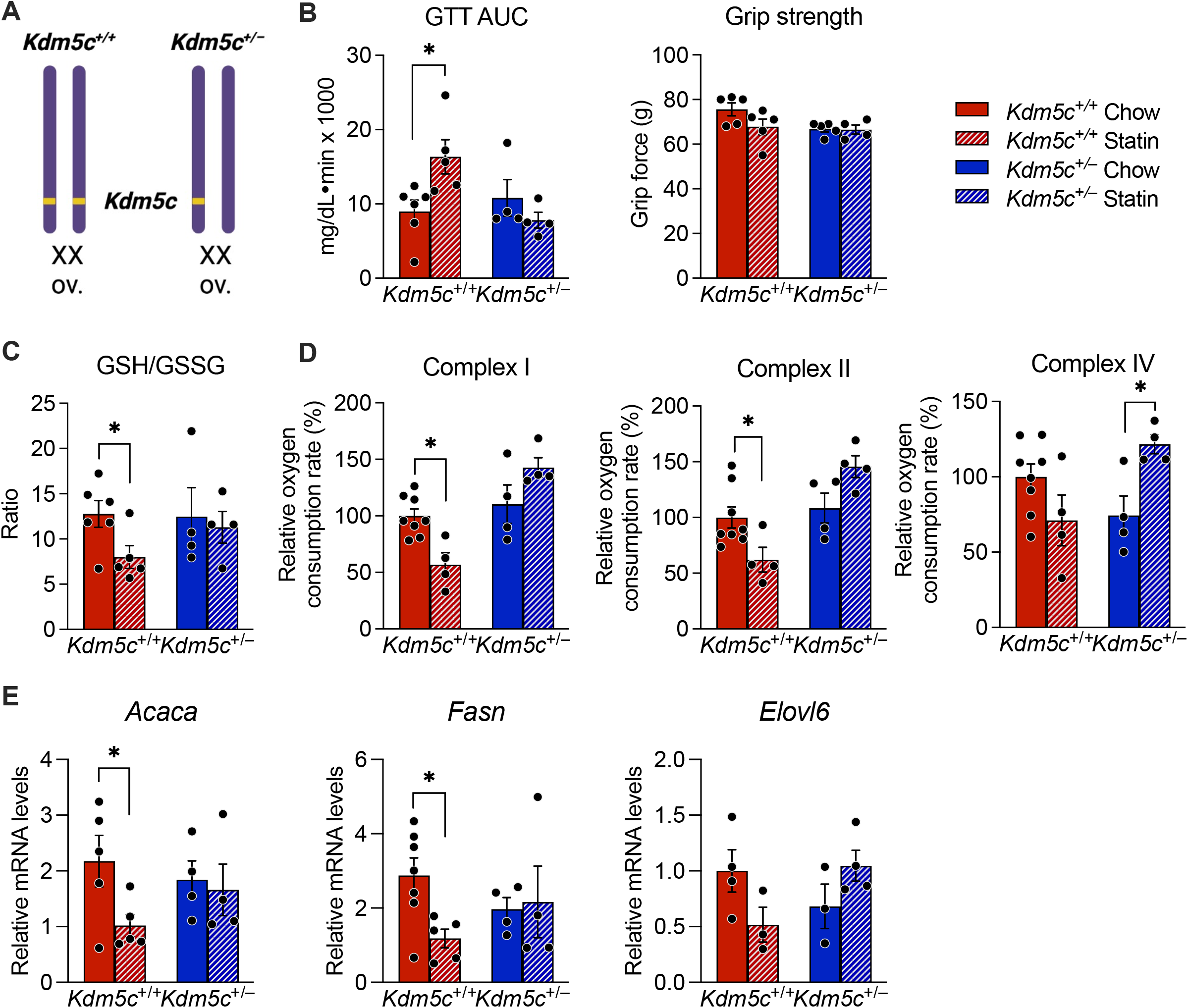
Female mice with *Kdm5c* gene dosage similar to males are protected from statin adverse effects. **(A)** Female XX mice carrying *Kdm5c* gene dosage similar to typical female (*Kdm5c*^+/+^) or typical male mice (*Kdm5c*^+/–^) were characterized for statin response. After 4 weeks on statin, mice were assessed for **(B)** glucose tolerance area under the curve and grip strength, **(C)** ratio of reduced to oxidized glutathione (GSH/GSSG), **(D)** mitochondrial respiration activity in complex I, II, and IV, and **(E)** fatty acid synthesis and elongase gene expression (see text for full gene names). Values represent mean ± SEM. *, *p* < 0.05 for t-tests within genotype for control and statin treatments.

### Women are more susceptible than men to statin-induced decrements in DHA ω-3 fatty acid and mitochondrial respiration

Our studies in mice indicated that reduction in DHA levels in female mice are a key determinant of statin adverse effects, as complementation with ω-3 fatty acids prevented glucose intolerance, reduced oxidative tone and impaired mitochondrial respiration. To determine relevance to humans, we assessed men and women for alterations in ω-3 fatty acid levels (DHA and EPA) in response to short-term statin treatment. Blood was collected from men and women without known cardiovascular disease or type 2 diabetes before statin administration and following 10 weeks of daily statin treatment (atorvastatin, 40 mg) (Abbasi et al., 2021) (**Fig. 6A**). Women and men both decreased DHA levels in response to statin, but in women the levels were reduced by twice as much, dropping more than 50 μmol/L compared to 25 μmol/L in men (**Fig. 6B**). Statin treatment cause a slight decrease in EPA levels in both sexes. Only women showed an increase in glucose levels after 10 weeks of statin treatment (Fig. 6B).

**Figure 6.**
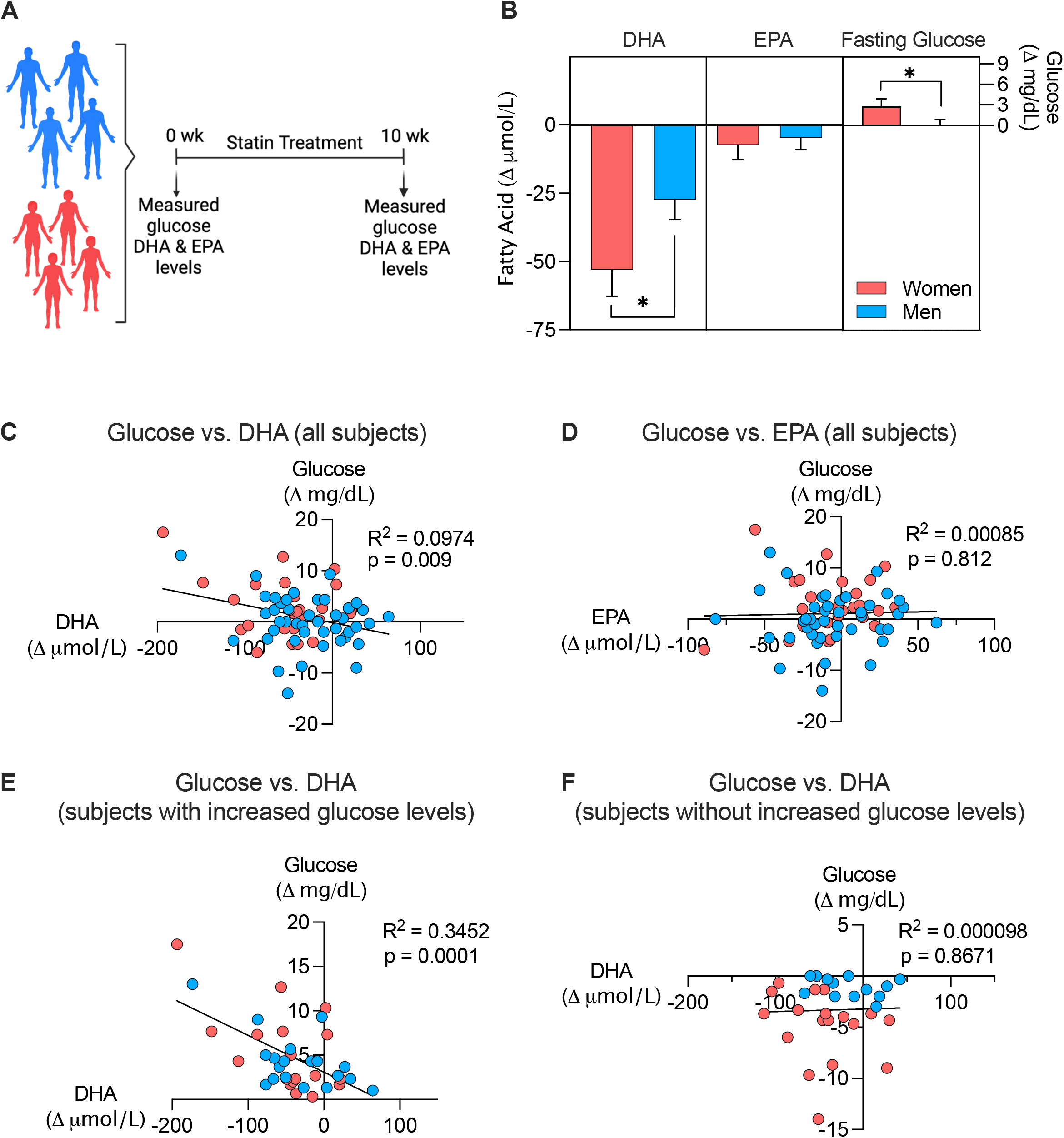
Women are more susceptible than men to statin-induced decrements in DHA levels. (A) Blood collected from statin-eligible men and women without type 2 diabetes was used to assess ω-3 fatty acid (DHA and EPA) and glucose levels before and at 10 weeks after statin treatment. **(B)** Women showed more pronounced statin-induced reduction in DHA levels, and increased glucose levels, compared to men. Data represent mean ± SEM; *, *p* < 0.05 in paired t-test between the sexes. **(C**,**D)** Regression of DHA or EPA levels vs. glucose levels in all subjects. **(E**,**F)** Regression of DHA or EPA vs. glucose levels for individuals that reduced fatty acid levels in response to statin treatment.

We further assessed the relationship between ω-3 fatty acid levels and alterations in glucose levels in response to short-term statin treatment. Analysis of all individuals in the study showed a weak but significant correlation between statin-induced decrement in DHA levels and increased glucose levels (**Fig. 6C**); no significant correlation was detected between EPA and glucose levels (**Fig. 6D**). Analysis of only individuals who experienced a decrease in DHA levels in response to statin revealed a stronger correlation between decrement in DHA and increased glucose levels (**Fig. 6E**), with no relationship between EPA and glucose levels (**Fig. 6F**).

To determine whether the mitochondrial deficiencies noted in statin-treated female mice are replicated in humans, we utilized patient-derived induced pluripotent stem cells (iPSC) from individuals who were susceptible or resistant to statin-induced NOD (**Fig. 7A**). Information was obtained from electronic medical records to identify statin NOD cases and statin-resistant controls. NOD cases had at least two of the following clinical indicators of diabetes arising during statin treatment: elevated fasting glucose levels (>126 mg/dL), diagnosis of diabetes during statin treatment, or first prescription of anti-diabetic drugs during statin treatment. Control subjects experience none of these indicators of NOD. Prior to statin initiation, NOD cases and controls both had normal fasting glucose measures (<110 mg/dL). iPSCs were derived from CD34+ peripheral blood mononuclear cells from male and female NOD cases and controls by reprogramming with Yamanaka factors; representative iPSC lines were validated for karyotype and expression of pluripotent markers (Kuang et al., 2019).

**Figure 7.**
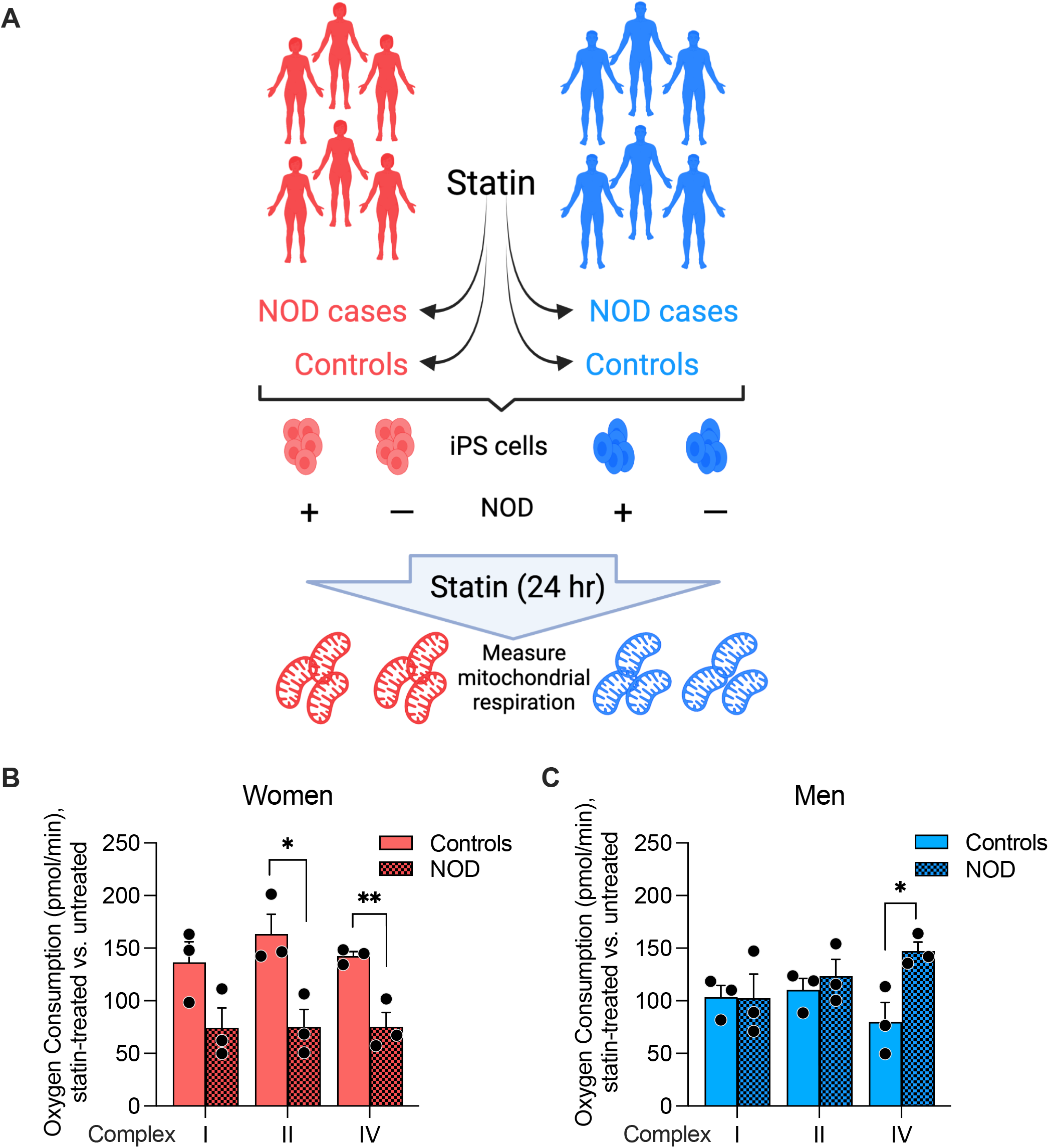
Impaired mitochondrial respiration in patient-derived iPSCs from women with statin new-onset diabetes (NOD). **(A)** iPSCs were programmed from peripheral blood mononuclear cells obtained from men and women with documented diabetes development during continuous statin (NOD cases) use or resistance to NOD (controls). iPSCs were cultured without or with statin for 24 hr, and frozen. Mitochondrial respirometry was performed on all samples in parallel using methodology for frozen samples (see Methods). Mitochondrial complex I, II and IV respiration in **(B)** female and **(C)** male NOD cases and controls. Values represent mean ± SEM. *, *p* < 0.05; **, *p* < 0.01 in t-test between control and NOD individuals.

We treated cultured iPSC lines (n = 3 independent lines per sex and disease status) with statin for 24 hr, and performed respirometry on all cell lines simultaneously. We utilized atorvastatin for these studies as we have determined that it produces a robust transcriptional response of genes in the mevalonate pathway in cultured iPSCs. We assessed the statin response of each patient cell line as the ratio of respiration in cells treated with statin compared to the same cells without statin treatment. Statin treatment impaired respiration in cells from women NOD cases compared to controls (**Fig. 7B**), consistent with our observations in the mouse. This effect was not observed in cell lines from men (**Fig. 7C**), suggesting potential differences in mechanisms underlying statin-associated NOD in women and men.

## Discussion

Statins remain the most commonly used drug to lower plasma cholesterol levels and reduce the risk of coronary heart disease. Adverse effects experienced by some statin users are a deterrent to continued use, leading to their reduced protection from cardiovascular disease. Previous studies have identified genetic variants that influence susceptibility to statin adverse effects in genes encoding the target of statin action (*HMGCR*, HMG-CoA reductase), a statin transporter (*SLCO1B1* organic anion transporter), and enzymes that metabolize statin for elimination (*ABCB1* transporter and *CYP3A* and *CYP2D6* cytochrome P450 genes) (Carr et al., 2019; Needham and Mastaglia, 2014; SEARCH Collaborative Group et al., 2008). Previous work has also implicated statin-induced reductions in the levels of lipid intermediates in the HMG-CoA pathway, including the mitochondrial electron transport component, coenzyme Q, or isoprenoid groups that form covalent modifications of numerous proteins (*e*.*g*., farnesol and geranylgeraniol) (Thompson et al., 2016; Ward et al., 2019). However, these genetic variants and biochemical mechanisms have not been related to differences in statin adverse effects between males and females.

We exploited multiple mouse models and human cohorts to assess sex differences in statin adverse effects and to identify molecular and genetic mechanisms. Our findings indicate that female C57BL/6J mice have impaired glucose tolerance and muscle grip strength after short-term statin treatment, whereas male mice on the same genetic background resist these adverse effects. Our findings further revealed that female sex interacts with statin treatment to reduce hepatic levels of the ω-3 fatty acid DHA, reduce cellular redox tone, and impair mitochondrial respiration. The central role of DHA was supported by the ability to prevent both the adverse biochemical and physiological effects of statin treatment in female mice by supplementation with fish oil. Our studies further determined that enhanced *Kdm5c* gene dosage in females (owing to expression from both X chromosomes in XX cells) is a determinant of both the biochemical and physiological adverse effects of statin. The role of X chromosome dosage as a determinant in sex differences has been shown for conditions such as obesity, autoimmunity, and neurodegeneration (Davis et al., 2020; Itoh et al., 2019; Link et al., 2020); the current studies extend the impact of X chromosome dosage to include response to statin.

Our results in the mouse were reinforced by findings that women experience more severe reductions than men in DHA levels after short-term (10 week) statin administration, and that DHA reduction was correlated with increases in fasting glucose levels. The specific reduction in DHA without reduction in the ω-3 fatty acid EPA that we observed is consistent with a previous study in a small Japanese cohort (Nozue and Michishita, 2015); however, results in that study were not stratified by sex. Additional translation of our findings in mouse to humans was achieved by demonstrating that cells derived from women, but not men, who developed NOD exhibited impaired mitochondrial function when treated with statin.

Our studies with mice and with cultured human cells allow us to assign biological sex distinct from gender as a primary component of increased female susceptibility to statin adverse effects. This is significant because gender may also influence the development and detection of NOD through effects on adherence to statin use, diet, physical activity, and likelihood of seeking medical attention for NOD. Additionally, we identified *Kdm5c* gene dosage as a component of biological sex that influences statin adverse effects. In humans as well as in mice, *KDM5C*/*Kdm5c* escapes X-chromosome inactivation to be expressed at higher levels in XX compared to XY tissues that are relevant to statin adverse effects, including liver, skeletal muscle, adipose tissue, and pancreas (Chen et al., 2012; Link et al., 2020). Here we explored a model of global reduction in *Kdm5c* gene dosage (*Kdm5c*^+/–^ female mice); future studies with tissue-specific modulation of *Kdm5c* dosage will be useful to zero in on the key tissue(s) that initiate statin NOD. Additionally, given the molecular function of KDM5C as a histone demethylase, future studies to identify KDM5C transcriptional targets in the presence of statin may identify additional genes that promote the development of NOD. Such findings may also be relevant to a better understanding of the development of type 2 diabetes in the general population.

In summary, our findings identify biochemical and genetic mechanisms for enhanced susceptibility of females to statin-related muscle weakness and impaired glucose homeostasis. They also suggest potential biomarkers for statin adverse effects (reduced DHA levels, glutathione levels, and mitochondrial function) and identify a novel candidate gene (*Kdm5c/KDM5C*) whose expression levels influence development of adverse effects. Importantly, our findings suggest a major role for altered ω-3 fatty acid metabolism in the pathogenesis of statin-induced NOD and myopathy and raise the possibility that supplementation with DHA may be of value in ameliorating or preventing these adverse effects.

## METHODS

### Mouse strains

Male and female C57BL/6J mice were obtained from The Jackson Laboratory (Bar Harbor, ME). C57BL/6 Four Core Genotypes mice with *Apoe* deficiency were described previously (AlSiraj et al., 2019) and were bred in-house as described for standard Four Core Genotypes mice (Link 2020). *Kdm5c*^+/+^ and *Kdm5c*^+/–^ were on a C57BL/6 background (floxed mice were originally backcrossed to C57BL/6J for 11 generations and then crossed to E2a-Cre transgenic mice) (Link et al., 2020). Male mice with global hemizygous *Kdm5c* deficiency (*Kdm5c*^− /Y^) were not recovered and thus only female mice were studied. Genotyping primers for Four Core Genotype *Apoe*^−/–^ and *Kdm5c* mice are provided in **Table S3**. Mice were maintained at ambient temperature on a 12h light/dark cycle. All animal studies were approved by the UCLA Institutional Animal Care and Use Committee.

### Administration of diets, oil gavage, and physiological measurements in live mice

At 8–10 weeks of age, mice were maintained on mouse chow (diet D1001 containing 10 kcal% fat, 20 kcal% protein and 70 kcal% carbohydrate; Research Diets, New Brunswick, NJ) or chow containing simvastatin (0.1 g/Kg body weight in mouse chow, formulation D11060903i, Research Diets) similar to previous reports (Muraki et al., 2012; Yokoyama et al., 2007). A simvastatin dose of 0.1g/Kg in mouse is similar to 80 mg/day in humans based on the FDA conversion factor for “human equivalent dose” in a mouse of 12.3. Thus, 80 mg per 60 Kg human = 1.3 mg/Kg. A dose of 0.1 mg/g (*i*.*e*., 0.1 g/Kg) for a 25 g mouse that eats 4 grams per day = 0.4 mg/25g body weight = 16 mg/Kg. Finally, 16 mg/Kg/12.3 (FDA conversion factor) = 1.3 mg/Kg.

For studies of ω-3 fatty acid co-therapy, fish oil dose was determined by metabolic rate adjustment of a human high-dose ω-3 fatty acid regimen (Terpstrat al., 2001). Mice fed chow or statin diets for 5 weeks were gavaged with a 10 μL/g bolus of coconut oil vehicle (Natural Source, New York, NY) or mixture of fish oil (Nordic Naturals, Watsonville, CA) and coconut oil (1:4), resulting in supply of 250 mg/Kg of DHA and 372.5 mg/Kg of EPA. Oral gavage was performed with 20 gauge plastic feeding tubes (FTP-20-30, Instech Laboratories, Plymouth Meeting, PA) 5 days/week.

Glucose tolerance was performed after a 5 hr fast (0800–1300) by intraperitoneal glucose injection (2 g/Kg body weight) and blood samples collected via tail nick at 0, 30, 60, 120, and 180 min. Glucose determinations were made with an AlphaTRAK glucometer (Zoetis, Parsippany, NJ). Grip strength was measured by spring scale dynamometer (model 8261-M, Ohaus, Parsippany, NJ).

### RNA-sequencing and qPCR

RNA was isolated from snap-frozen liver samples with TRIzol (Thermo Fisher Scientific). Reverse transcription was performed with iScript and qPCR with SsoAdvanced SYBR Green Supermix (Bio-Rad, Hercules, CA) using primers in **Table S2**.

RNA-seq was performed on n=3 livers from female and male mice each fed chow or chow-statin diets. RNA-seq libraries were generated as we have previously (Link et al., 2020) with a workflow consisting of poly (A) RNA selection, RNA fragmentation, oligo(dT) priming and cDNA synthesis, adaptor ligation to double-stranded DNA, strand selection and PCR amplification to generate final libraries. Following quantitation (Qubit) and quality-check (4200 TapeStation (Agilent), index adaptors were used to multiplex samples. Sequencing was performed at the UCLA Technology Center for Genomics & Bioinformatics on a NovaSeq 600 sequencer to obtain 50 bp paired-end reads. RNA-seq alignment was performed as described (Link 2020). The average read depth was 18 × 10^6^ reads/sample, with 85% of reads aligning uniquely to the genome (GRCm38.97). Differential expression was identified by EdgeR (adjusted p<0.05) (Robinson et al., 2010). Pathway analysis for differentially expressed genes was performed with Enrichr (Xie et al., 2021). Volcano plots were generated with ggplot2 package (Wickham, 2016).

### Metabolomic analyses

Nonpolar lipid metabolites were quantified in mouse liver extracts as previously (Benjamin et al., 2013). Livers were extracted in chloroform:methanol:PBS with inclusion of internal standards C12:0 dodecylglycerol (10 nmol) and penta-decanoic acid (10 nmol). Organic and aqueous layers were separated, and the aqueous layer was acidified [for detection of metabolites such as lysophosphatidic acid (LPA) and LPA-ether (LPAe)], followed by re-extraction with chloroform. The organic layers were combined, dried down, and analyzed by both single-reaction monitoring (SRM)-based LC-MS/MS as described (Benjamin et al., 2013). Metabolites were quantified by integrating the area under the peak and were normalized to internal standard values and then levels from statin-treated animals were expressed as relative levels compared with control animals.

### Biochemical measurements

Plasma glutamate, α-ketoglutarate, and glutathione levels were assayed using enzymatic kits (Glutamate Assay Kit, Sigma-Aldrich, MAK004-1KT; α-Ketoglutarate Assay Kit, Sigma-Aldrich, MAK054-1KT; Glutathione Assay Kit, Cayman Chemical, #703002) according to the manufacturer’s instructions. Prior to glutamate or α-ketoglutarate assays, plasma was deproteinated using a Microcon-10kDa centrifugal filter unit (Millipore Sigma, #MRCPRT010). For glutathione (GSH) assays, plasma was deproteinated using MPA reagent (metaphosphoric acid, 1 g/ml, Sigma-Aldrich 239275) and TEAM reagent (triethanolamine, Sigma-Aldrich, #T58300). Direct assessment of deproteinated samples provided GSH levels. To assay levels of the oxidized form of glutathione (GSSG), GSH in deproteinated samples was derivatized with 2-vinylpyridine (Sigma-Aldrich, #13229-2) prior to assessment. To assay glutamate, α-ketoglutarate, and glutathione in liver, homogenates were prepared in ice-cold buffer specific for the component being analyzed. Buffer was provided in the Glutamate Assay Kit (for glutamate assessment), in the α-ketoglutarate assay kit (for α-KG assessment), or was 50mM 2-(N-morpholino)ethanesulfonic acid (MES) buffer (pH6) with 1mM EDTA (for glutathione) (Baechler et al., 2019; Edwards et al., 2021; Jia et al., 2010). Samples were deproteinated and assayed as described above for plasma samples. Plasma lipid levels were determined by enzymatic assays as previously described (Link et al., 2015). Plasma insulin levels were determined with a Mouse Ultrasensitive Insulin ELISA kit (ALPCO, Salem, NH).

### Mitochondrial respiration assays

Mitochondrial respirometry was performed using homogenates of frozen liver with cytochrome C reconstitution as described (Acin-Perez et al., 2020; Ngo et al., 2021). This method allows the assessment of complex I, II and IV activities. Briefly, homogenates were prepared in MAS buffer (70 mM sucrose, 220 mM mannitol, 5 mM KH2PO4, 5 mMgCl2, 1 mM EGTA, 2 mM HEPES, pH7.4) in a Dounce homogenizer, cleared by centrifugation (1000g for 10 min, 4°C), and the final supernatant was collected and protein quantified with Bio-Rad Protein Assay (Hercules, CA). Respirometry studies to detect Complex I, Complex II, and Complex IV activity were performed with a Seahorse XF96 analyzer in MAS buffer containing 10 μg/ml cytochrome C. Substrate injections were: Port A)—pyruvate + malate (5 mM each) for Complex I, 5 mM succinate (Complex II), or 5 mM rotenone (Complex IV); Port B— 2 μM rotenone + 4 μM antimycin; Port C—0.5 mM TMPD (tetramethyl phenylene diamine) + 1 mM ascorbic acid; and Port D—50 mM azide.

### Human blood samples

Samples assessed here were collected for a previously published study (Abbasi et al., 2021). Briefly, volunteers who were eligible for statin therapy for cardiovascular disease prevention and did not have type 2 diabetes, statin intolerance, or other exclusion criteria [detailed in (Abbasi et al., 2021)] provided written informed consent. Data from these subjects was previously published demonstrating that atorvastatin treatment for 10 weeks increased insulin resistance in individuals without type 2 diabetes (Abbasi et al., 2021), but no assessments of ω-3 fatty acids nor sex stratification were performed in the previous study. For data presented here, ω-3 fatty acids DHA and EPA were assessed in blood samples obtained at week 0 and at week 10 of atorvastatin treatment in 70 subjects (44 men and 26 women). DHA and EPA levels were determined using the Serum Comprehensive Fatty Acid Panel (C8–C26), which employs gas chromatography/tandem mass spectrometry (Quest Diagnostic Nichols Institute, San Juan Capistrano, CA). Glucose levels in these samples were previously determined (Abbasi et al., 2021).

### Patient-derived iPSCs

We used electronic health records from Kaiser Permanente of Northern California (KPNC) to identify statin users who developed new-onset diabetes (NOD). NOD cases were defined as those who had his/her first statin (simvastatin, lovastatin, atorvastatin, pravastatin, rosuvastatin, or pitavastatin; **Table S3**) prescription at age 40–75, and documented continuous statin use for 3 years. Continuous statin use was defined as having greater than 8 30-day prescription refills per year or greater than 3 90-day prescription refills per year. Individuals prescribed statin combinations were excluded.

Individuals with evidence of diabetes prior to the date of the first statin prescription or within the first 3 months after the start of statin use were excluded. Evidence of diabetes included either i) diabetes diagnosis based on ICD9 codes for type 1, 2 or gestational diabetes (**Table S4** or ii) prescription for glucose lowering drugs (alpha-glucosidase inhibitors, dipeptidyl peptidase-4 inhibitors, meglitinides, sulfonylureas, biguanides, and thiazolidinediones) and insulin (all formulations shown on **Table S5**). In addition, individuals with prescriptions for glucose raising drugs, such as oral corticosteroids (**Table S6**) or with ICD9/CPT codes for bariatric surgery (**Table S7**), in the 3 years prior to start of statin treatment, through the 3 years after the start of statin treatment were excluded.

To ensure that individuals were not diabetic prior to the start of statin treatment, individuals were required to have at least two fasting glucose (FG) measures of 50–110 mg/dL and within 30 mg/dL of each other. Individuals with an outpatient FG measure ≥ 126 mg/dL within the first 3 months after the start of statin treatment were excluded. FG values < 30 mg/dL or > 600 mg/dL were also excluded. Individuals who developed NOD in the 3 years after statin initiation were identified and recruited, with diabetes defined as two or more of the following diagnostic elements: 1) FG ≥ 126 mg/dL, 2) diabetes ICD9 code, and/or 3) prescription of a glucose lowering drug (Table S5).

Controls were defined as statin users with normal glycemia (all FG values of 50–110 mg/dL) in the 3 years prior to statin initiation and during the first 5 years on-treatment with evidence of continuous statin prescription. Individuals with a diabetes ICD code or use of glucose-modifying drug during the 8 year window were excluded. Blood samples were drawn from identified NOD cases and controls, with clinical and demographic information shown in **Table S8**. Written informed consent was obtained from all study subjects and studies were performed with approval from the Institutional Review Boards at Kaiser Permanente Northern California and the UCSF Benioff Children’s Hospitals.

Patient-derived iPSCs were prepared from the participants (**Table S8)**. Briefly, iPSCs were reprogrammed from CD34^+^ peripheral blood mononuclear cells (PBMCs), and authenticated as previously described (Kuang et al., 2019). Cells were cultured in mTESR1 media at 37°C at 5% CO2 and passaged using accutase (Stemcell Technologies, Cat. # 07920) and media supplemented with Y-27632 2HCl (ROCK) inhibitor (Selleckchem, Cat. # S1049; 10μM working stock). Cells were incubated with either 250μM atorvastatin (Sigma-Aldrich #189291) dissolved in ethanol or mock buffer (ethanol only) for 24 hours, after which cells were washed with PBS and flash frozen. Mitochondrial respiration was determined as described (Acin-Perez et al., 2020; Ngo et al., 2021) and detailed under “Mitochondrial respiration assays.”

### Availability of data and biological materials

RNA-seq data associated with Fig. 2 have been deposited in GEO (Accession number GSE184588). We will provide mouse strains described in this study upon request to non-commercial entities for research purposes in agreement with University of California regulations.

### Statistics

Statistical tests used for each figure are indicated in legends using Prism (GraphPad Software, San Diego, CA). In general, in studies of mice on chow vs. statin diets, within sex group comparisons were performed with Student’s *t* test (two-tailed). For analysis of Four Core Genotypes, groups were compared with three-factor ANOVA with main factors of sex chromosome complement (XX vs. XY), gonadal sex (ovaries vs. testes), and treatment (chow vs. statin). For analysis of fish oil treatment, groups were compared with two-factor ANOVA with main factors of diet (chow vs. statin) and co-therapy (coconut oil vs. fish oil). When two-way or three-way ANOVA analyses were significant, subsequent relevant pairwise analyses were performed with Student’s *t* test. Error bars represent SEM.

## Acknowledgments

The authors thank Katherine Scheker, Lara Bideyan, and Carlos Galvan for assistance with biochemical assays or data analysis in early stages of this study. We thank Elizabeth Theusch for assembling tables related to human patient-derived iPSC lines. We thank the POST study subjects, as well as Brendan Neilan and Gilbert Nalula for their assistance in generation of the iPSC lines. These studies were supported by P50 GM115318 from the National Institute of General Medical Sciences (RMK, MWM, KR), U54 DK120342 from the National Institute of Diabetes and Digestive and Kidney Disease/Office of Research on Women’s Health (KR), R21 AR077782 (KR and PZ), The Iris Cantor-UCLA Women’s Health Center/National Center for Advancing Translational Sciences and UCLA Clinical and Translational Science Institute (KR), NIH pre-doctoral fellowships F31 F31AA028183 (JJM) and T32DA024635 (JJM), American Heart Association post-doctoral fellowship 20POST35100000 (CBW), and Doris Duke Foundation, R01 DK116750, P30 DK116074, and R01 DK120565 (JWK).

## Author Contributions

PZ, JJM, and KR designed studies and wrote the manuscript. PZ, JJM, and ER performed mouse studies. LV performed respirometry studies. PZ analyzed transcriptome and lipodome data, and CBW analyzed human fatty acid data. PZ and CBW prepared figures. JCL generated mouse cohorts and aligned RNA-seq data. DKN provided lipidomic measurements. CI, ML, G S-K, and A O-O analyzed electronic medical records and identified NOD and control subjects. FA and JWK provided human blood samples for ω-3 fatty acid analyses; MJM performed ω-3 fatty analyses. RK advised on statin studies and edited the manuscript. MWM, AM, and YLK derived and cultured iPSCs. All authors reviewed the manuscript.

## Supplemental Figures

**Suppl. Figure 1.**
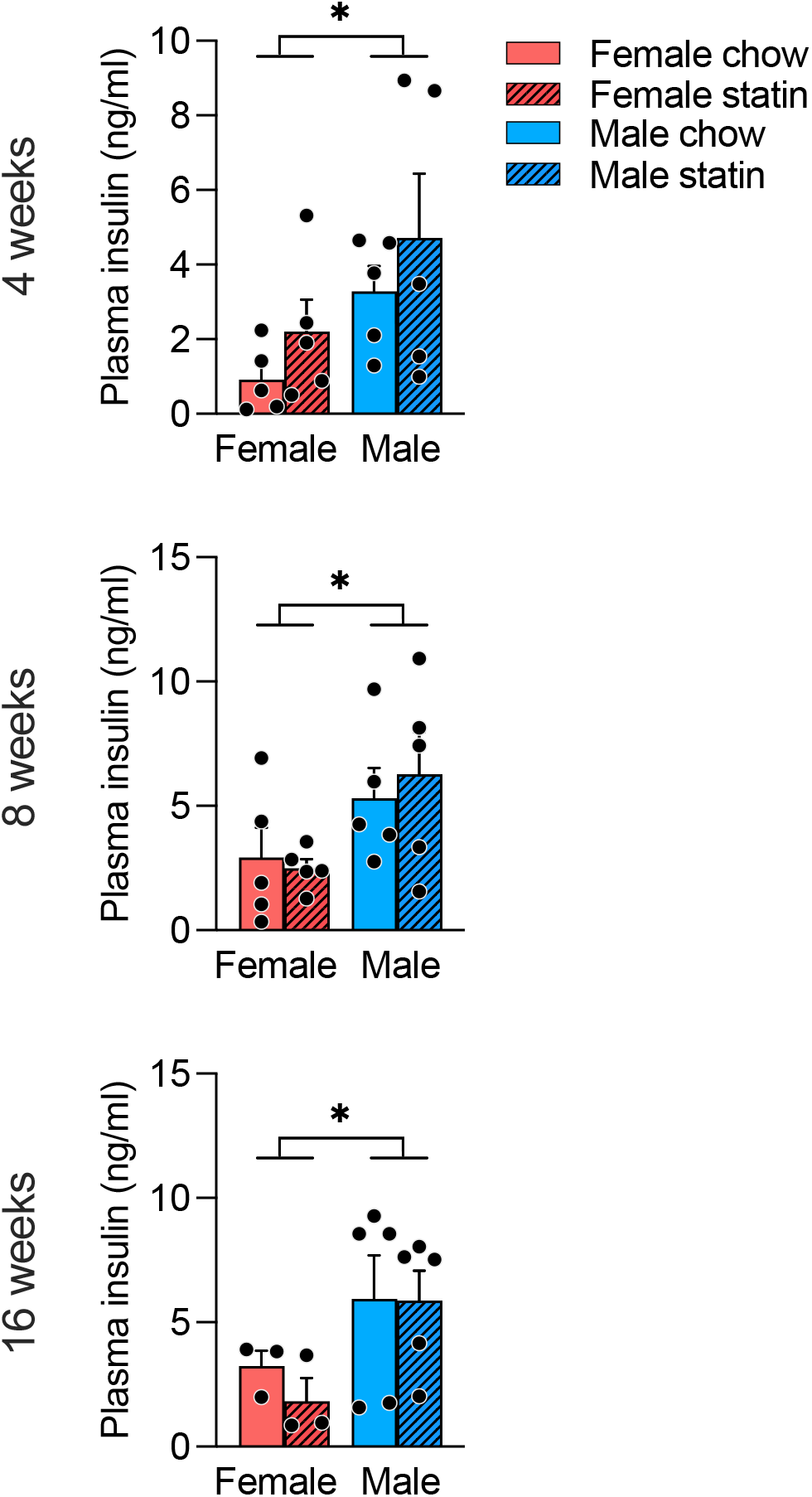
Effects of statin on plasma insulin concentrations in male and female mice. Groups were compared by 2-way ANOVA with main factors of diet (chow vs. statin) and sex (female vs. male). Values represent mean ± SEM. *, *p* < 0.05.

**Suppl. Figure 2.**
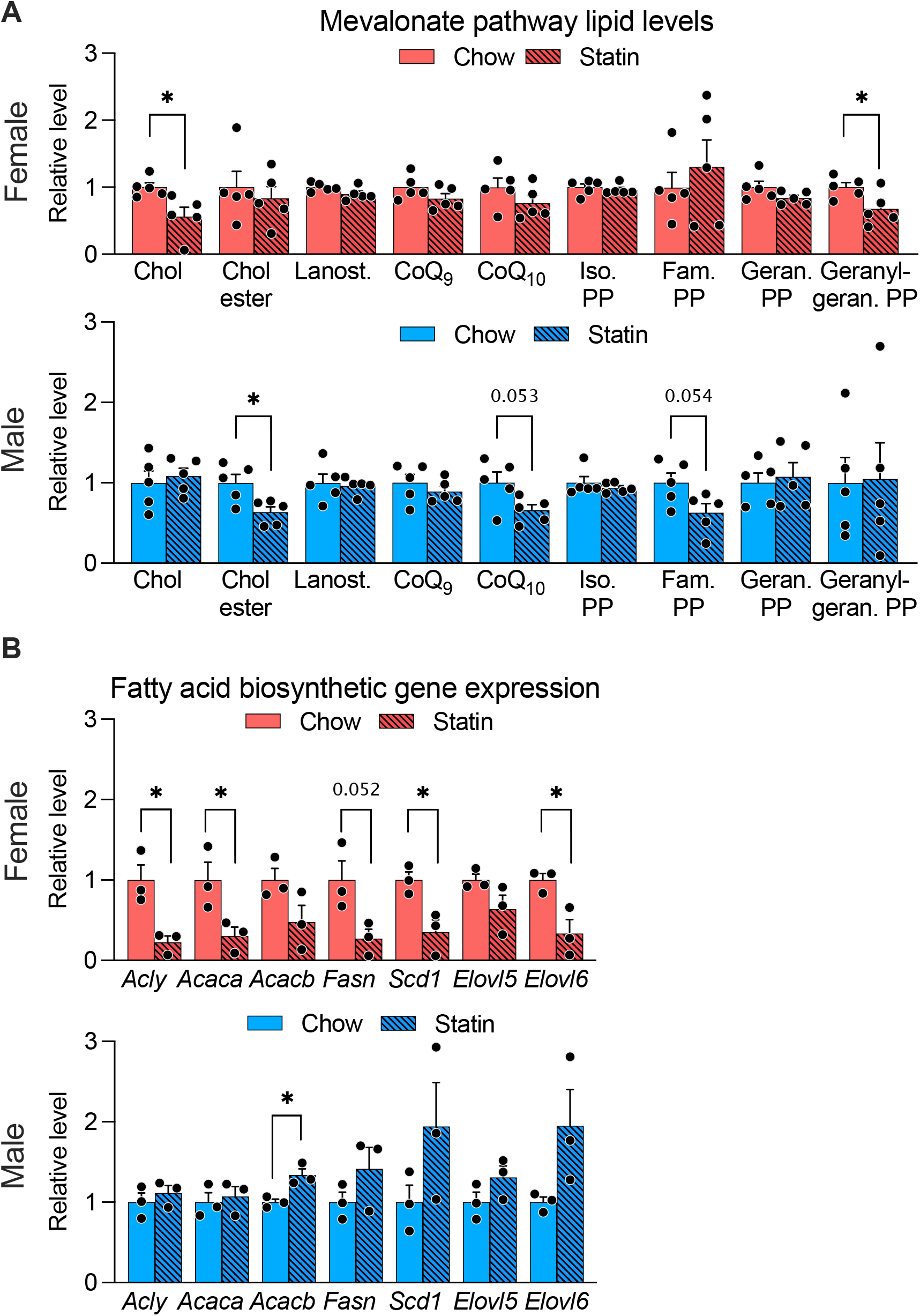
Sex differences in statin effects on cholesterol pathway intermediates and fatty acid biosynthetic gene expression. (A) Statin-induced alterations in hepatic lipids within the mevalonate pathway in male and female mice. Chol, cholesterol; Lanost., lanosterol; CoQ, coenzyme Q; Iso PP, isopentenyl pyrophosphate; Farn. PP, farnesyl pyrophosphate; Geran. PP, geranyl pyrophosphate; Geranyl geran. PP, geranylgeranyl pyrophosphate. (B) Statin-induced alterations in hepatic gene expression for fatty acid synthetic genes. *Acly*, ATC citrate lyase; *Acaca*, acetyl-CoA carboxylase α; *Acacb*, acetyl-CoA carboxylase β *Fasn*, fatty acid synthase; *Scd1*, stearoyl-CoA desaturase 1; *Elovl5*, fatty acid elongase 5; *Elovl6*, fatty acid elongase 6. Brackets show significant paired t-tests with each sex for a specific lipid or mRNA; *, *p* < 0.05.

