## Supplemental Table 2 for "X chromosome dosage drives statin-induced dysglycemia and mitochondrial dysfunction"

Gene expression qPCR primers

| | |
|---|---|
| *Gclc* (forward) | 5'- TGCACATCTACCACGCAGTCAA -3' |
| *Gclc* (reverse) | 5'- TCAAGAACATCGCCTCCATTCA -3' |
| *Glud1* (forward) | 5'- GCAACCATGTGTTGAGCCTCT -3' |
| *Glud1* (reverse) | 5'- CCACAGCGCACTTGTATGTCA -3' |
| *Got1* (forward) | 5'- CGCCTAGTTCTTGGGGACAAC-3' |
| *Got1* (reverse) | 5'- TCCCAGGTTGGTGATGATACG -3' |
| *Fasn* (forward) | 5'- GTTGGCCCAGAACTCCTGTA -3' |
| *Fasn* (reverse) | 5'- GTCGTCTGCCTCCAGAGC -3' |
| *Acaca* (forward) | 5'- GCCTCTTCCTGACAAACGAG -3' |
| *Acaca* (reverse) | 5'- TGACTGCCGAAACATCTCTG -3' |
| *Elovl6* (forward) | 5'- GATGACCAAAGGCCTGAAGC -3' |
| *Elovl6* (reverse) | 5'- GTGGTGGTACCAGTGCAGGA -3' |

Genotyping primers

| | |
|---|---|
| *Tg Sry* (forward) | 5'- AGCCCTACAGCCACATGATA -3' |
| *Tg Sry* (reverse) | 5'- GTCTTGCCTGTATGTGATGG -3' |
| *Ymt* (forward) | 5'- CTGGAGCTCTACAGTGATGA -3' |
| *Ymt* (reverse) | 5'- CAGTTACCAATCAACACATCAC -3' |
| *Myo* (forward) | 5'- TTACGTCCATCGTGGACAGCAT -3' |
| *Myo* (reverse) | 5'- TGGGCTGGGTGTTAGTCTTAT -3' |
| oIMR0180 | 5'- GCCTAGCCGAGGGAGAGCCG -3' |
| oIMR0181 | 5'- TGTGACTTGGGAGCTCTGCAGC -3' |
| oIMR0182 | 5'- GCCGCCCCGACTGCATCT -3' |
